# Superior colliculus projections drive dopamine neuron activity and movement but not value

**DOI:** 10.1101/2024.10.04.616744

**Authors:** Carli L. Poisson, Izzabella K. Green, Gretchen M. Stemmler, Amy R. Wolff, Julianna Prohofsky, Cassandra Herubin, Madelyn Blake, Benjamin T. Saunders

## Abstract

To navigate dynamic environments, animals must rapidly integrate sensory information and respond appropriately to gather rewards and avoid threats. It is well established that dopamine (DA) neurons in the ventral tegmental area (VTA) and substantia nigra (SNc) are key for creating associations between environmental stimuli (i.e., cues) and the outcomes they predict. Critically, it remains unclear how sensory information is integrated into dopamine pathways. The superior colliculus (SC) receives direct visual input and is positioned as a relay for dopamine neuron augmentation. We characterized the anatomical organization and functional impact of SC projections to the VTA and SNc in rats. First, we show that neurons in the deep layers of SC synapse densely throughout the ventral midbrain, interfacing with projections to the striatum and ventral pallidum, and these SC projections excite dopamine and GABA neurons in the VTA/SNc in vivo. Despite this, cues predicting SC→VTA/SNc neuron activation did not reliably evoke behavior in an optogenetic Pavlovian conditioning paradigm, and activation of SC→VTA/SNc neurons did not support primary reinforcement or produce place preference/avoidance. Instead, we find that stimulation of SC→VTA/SNc neurons evokes head turning. Focusing optogenetic activation solely onto dopamine neurons that receive input from the SC was sufficient to invigorate turning, but not reinforcement. Turning intensity increased with repeated stimulations, suggesting that this circuit may underlie sensorimotor learning for exploration and attentional switching. Together our results show that collicular neurons contribute to cue-guided behaviors by controlling pose adjustments through interaction with dopamine neurons that preferentially engage movement instead of reward.

## INTRODUCTION

To make sense of dynamic environments, animals rely on sensory information (i.e., cues) that, through Pavlovian learning, comes to predict biologically-relevant events, including rewards and threats. It has been well established that the midbrain dopamine system is crucial for the creation (Handler et al., 2019; Tsai et al., 2009) and expression (Fischbach & Janak, 2019; van Zessen et al., 2021) of cue-evoked conditioned behaviors (Berke, 2018; Berridge & Robinson, 1998; Mohebi et al., 2019; Saunders et al., 2018; Sharpe et al., 2017; Wise, 2009; Wise & Jordan, 2021). Notably, dopamine neurons respond to visual cues at faster latencies than would be possible through the canonical visual cortical processing pathway (da Silva et al., 2018; Redgrave et al., 1999; Schultz, 2007), suggesting a direct route of sensory information transmission to dopaminergic regions (Redgrave & Gurney, 2006). Despite this, the mechanisms underlying sensory integration with dopamine circuits remain poorly understood, and inputs providing rapid sensory augmentation have not been well characterized.

One candidate for providing direct sensory information is the superior colliculus (SC), as it receives direct retinal input and is key for navigating space (Ito & Feldheim, 2018; May, 2006) and shifting attention (Dean et al., 1989; Krauzlis et al., 2013). The SC has been shown to influence behavior via encoding of choice (Steinmetz et al., 2019; Thomas et al., 2023) and rewards (Ikeda & Hikosaka, 2007). The intermediate and deep layers of the SC are thought to act as a saliency gate, filtering and routing the most relevant information about the environment to downstream regions (Basso et al., 2021; Bertram et al., 2014; Bromberg-Martin et al., 2010a; Cooper et al., 1998; Evans et al., 2018; Lovejoy & Krauzlis, 2010; Mysore & Knudsen, 2011; Redgrave & Gurney, 2006; Wang et al., 2020; White et al., 2017; Lee et al., 2020), and some neurons there directly project to neurons in the VTA and SNc (Coizet et al., 2003; Comoli et al., 2003; Dawbarn & Pycock, 1982; May et al., 2009; McHaffie et al., 2006; Pradel et al., 2025). In primates and rodents, SC is active during reward-related tasks (Viviani et al., 2020) and involved in appetitive Pavlovian conditioning to visual cues (Takakuwa et al., 2017). In anesthetized rats, visual stimuli activate deep layer SC neurons, which is necessary for rapid dopamine neuron activity modulation in the ventral midbrain (Comoli et al., 2003; Redgrave et al., 2010). Further, disinhibition of the SC, but not visual cortex, allows for dopamine neurons to be responsive to visual stimuli in anesthetized rodents (Dommett et al., 2005). Two recent studies suggest that SC neurons projecting specifically to the ventral midbrain are important for directing attention in appetitive environments by controlling head orientation during hunting and social interactions (Huang et al., 2021; Solié et al., 2022).

Thus, the neural architecture for routing of sensory information to dopamine neurons via the SC is in place, but how this circuit functions in the context of cue-based learning, value assignment, and attention remains unclear (Bromberg-Martin et al., 2010a; Kaźmierczak & Nicola, 2022; Redgrave & Gurney, 2006). Here, we investigated how SC projections to the VTA/SNc impact dopamine neuron activity and subsequent motivated behaviors in awake, freely behaving rats. Using a combination of approaches, we find that SC neurons projecting to the ventral midbrain activate dopamine neurons and drive postural changes without creating conditioned behavior or producing strong valence representations. Our results highlight a brain circuit that is important for guiding movement to redirect attention, via interaction with classic learning systems, and suggest that distinct subpopulations of dopamine neurons are preferentially engaged to influence movement versus reward.

## RESULTS

### Superior colliculus projections to the VTA and SNc

We first examined superior colliculus (SC) projections to the rat ventral midbrain, using three complementary approaches (**Fig 1**). Injection of an AAV coding for GFP into the SC resulted in general expression throughout the intermediate and deep layers. Terminals from these neurons were visible throughout the ventral midbrain, including the VTAand SNc (**Fig 1A,C**). We counterstained this tissue for TH, demonstrating intermingling of SC neuron fibers with dopamine neurons in both regions (**Fig 1C**). To verify that these SC projections make synapses in the VTA/SNc, we injected the SC with a virus coding for the expression of membrane-bound GFP and mRuby conjugated to synaptophysin (**Fig 1D**), a synaptic labeling protein (Beier et al., 2015). This resulted in strong GFP expression in the VTA/SNc, reflecting axon terminal membranes, adjacent to dense mRuby puncta, indicative of synaptic connections with VTA/SNc neurons (**Fig 1E-G**). Finally, we examined the projection patterns of VTA/SNc neurons that receive monosynaptic input from neurons in the SC. To do this, we made use of the anterograde transsynaptic transport properties of AAV1 (Zingg et al., 2017). An AAV1 coding for cre recombinase was injected into the SC, along with the injection of a second AAV coding for cre-dependent expression of mCherry into the VTA or SNc (**Fig 1H**). This resulted in strong mCherry expression in neurons in the VTA and SNc (**Fig 1I**). Examination of forebrain regions showed strong mCherry expressing terminal fibers throughout the dorsal and ventral striatum and ventral pallidum (**Fig 1J**). Together these results build on previous findings (Huang et al., 2021; Redgrave, Coizet, Comoli, McHaffie, et al., 2010; Solié et al., 2022), demonstrating strong innervation of VTA/SNc by the SC, including to neurons projecting to the broader meso-striatal network.

**Fig 1.**
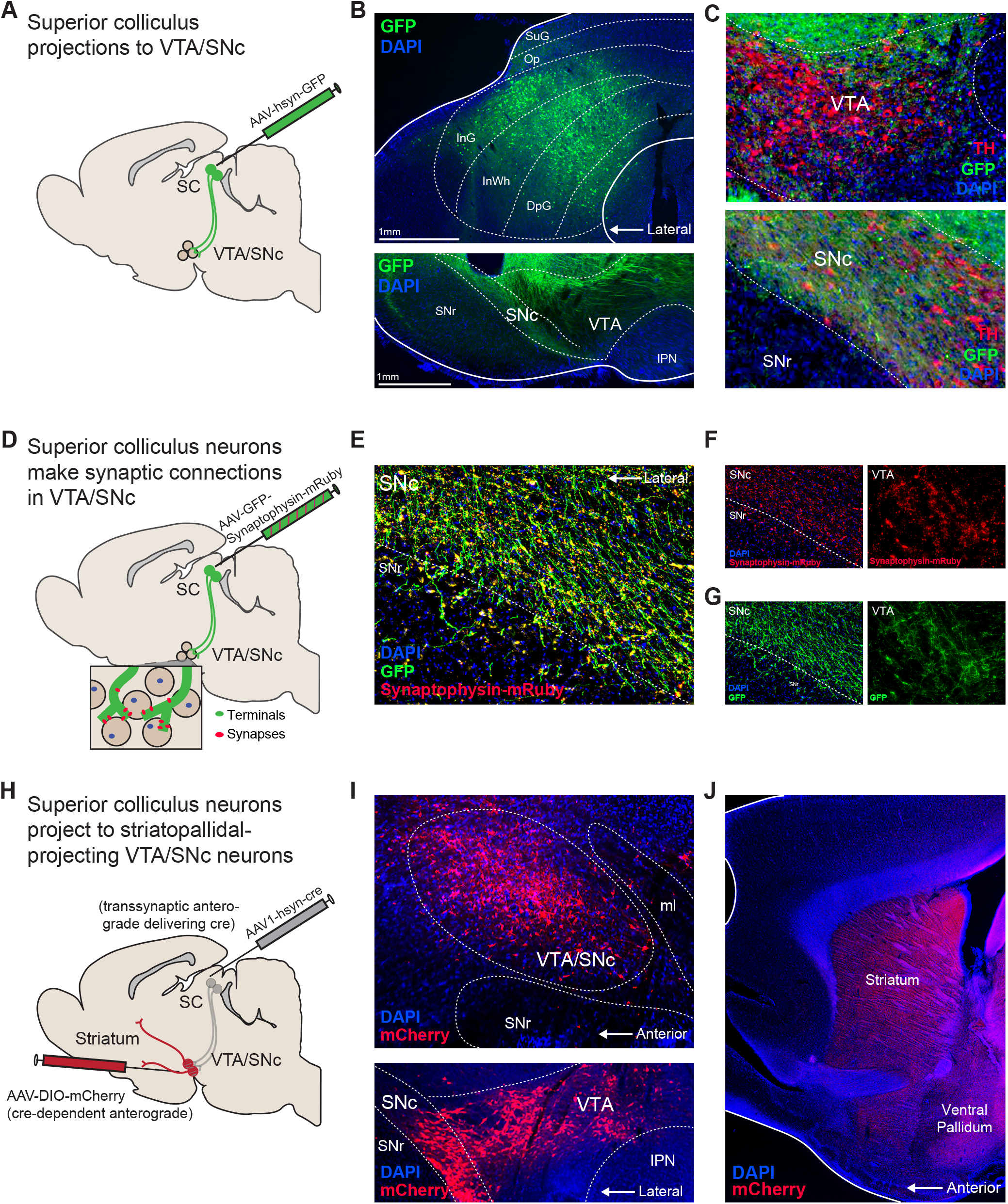
Superior colliculus projections to the ventral midbrain. A) Viral approach for targeting SC neurons. B) Injection of a GFP-expressing virus (green) into the SC resulted in expression throughout the intermediate and deep layers, with terminals visible throughout the ventral midbrain in the VTA and SNc. C) Tissue was counterstained for tyrosine hydroxylase (TH, red), demonstrating dense intermingling of SC projections with DA neurons in the VTA and SNc. D) Viral approach for visualizing monosynaptic connections between SC and the ventral midbrain. E) Injection of a virus coding for membrane-bound GFP and mRuby conjugated to the synaptophysin protein in the SC demonstrated strong innervation of the VTA and SNc. F) Dense mRuby puncta, indicating synaptic contacts between SC terminals and ventral midbrain neurons, were seen in VTA and SNc, G) closely associated with GFP-expressing terminals. H) Viral approach for transsynaptic tracing of SC-forebrain circuits. An AAV1 virus delivering cre recombinase was injected into the SC, combined with injection of a cre-dependent virus coding for mCherry into the VTA/SNc. I) Via the anterograde transsynaptic transport of AAV1-cre, we visualized VTA/SNc neurons receiving monosynaptic inputs from the SC (top: sagittal view; bottom: coronal view). J) mCherry fibers from VTA/SNc neurons receiving SC input were evident throughout the striatum and ventral pallidum in the forebrain.

### Superior colliculus projections excite VTA/SNc dopamine neurons *in vivo*

Above we demonstrate that SC neurons project strongly to the VTA and SNc. These inputs are thought to be largely glutamatergic (Redgrave, Coizet, Comoli, McHaffie, et al., 2010; Z. Zhou et al., 2019), but the general impact of SC neuron activity on the ventral midbrain neurons *in vivo* is unclear. To explore this, we used a combination of fiber photometry and optogenetics. GCaMP8f was expressed in dopamine neurons in TH-cre rats, to record population-level dopamine neuron activity in the VTA and SNc (**Fig 2A,B**) during optogenetic activation of SC terminals expressing the red-shifted opsin ChrimsonR. We delivered brief unsignaled laser stimulation at 5 or 20 Hz for 5-sec intervals. Examining video recordings of these laser delivery windows, we found that laser evoked movement invigoration, which scaled in intensity at the 20-Hz frequency (**Fig. 2C**: paired t test, t(11)=3.27, p=.0075). Across stimulation sites, 5-Hz activation (**Fig 2D**) produced a robust increase in dopamine neuron activity (**Fig 2E,F**; one-sample t test peak, t(10)=5.37, p=.0003; AUC t(10)=6.165, p=.0001). VTA and SNc responses differed in signal peak (**Fig 2G**; unpaired t test t(9)=2.548, p=.0313), but not overall AUC (**Fig 2H**; t test t(9)=.7098, p=.496) during the stimulation window. SC terminal stimulation at 20 Hz (**Fig 2I**) also robustly increased dopamine neuron activity (**Fig 2J,K**; one sample t test peak t(10)=5.385, p=.0003; AUC t(10)=2.557, p=.0285). VTA and SNc responses at 20 Hz did not differ (**Fig 2Q,R**; t test for peak t(9)=.499, p=.6299; t test for AUC t(9)=.281, p=.785). Together these data show that SC terminals acutely activate dopamine neurons in the VTA and SNc *in vivo*.

**Fig 2.**
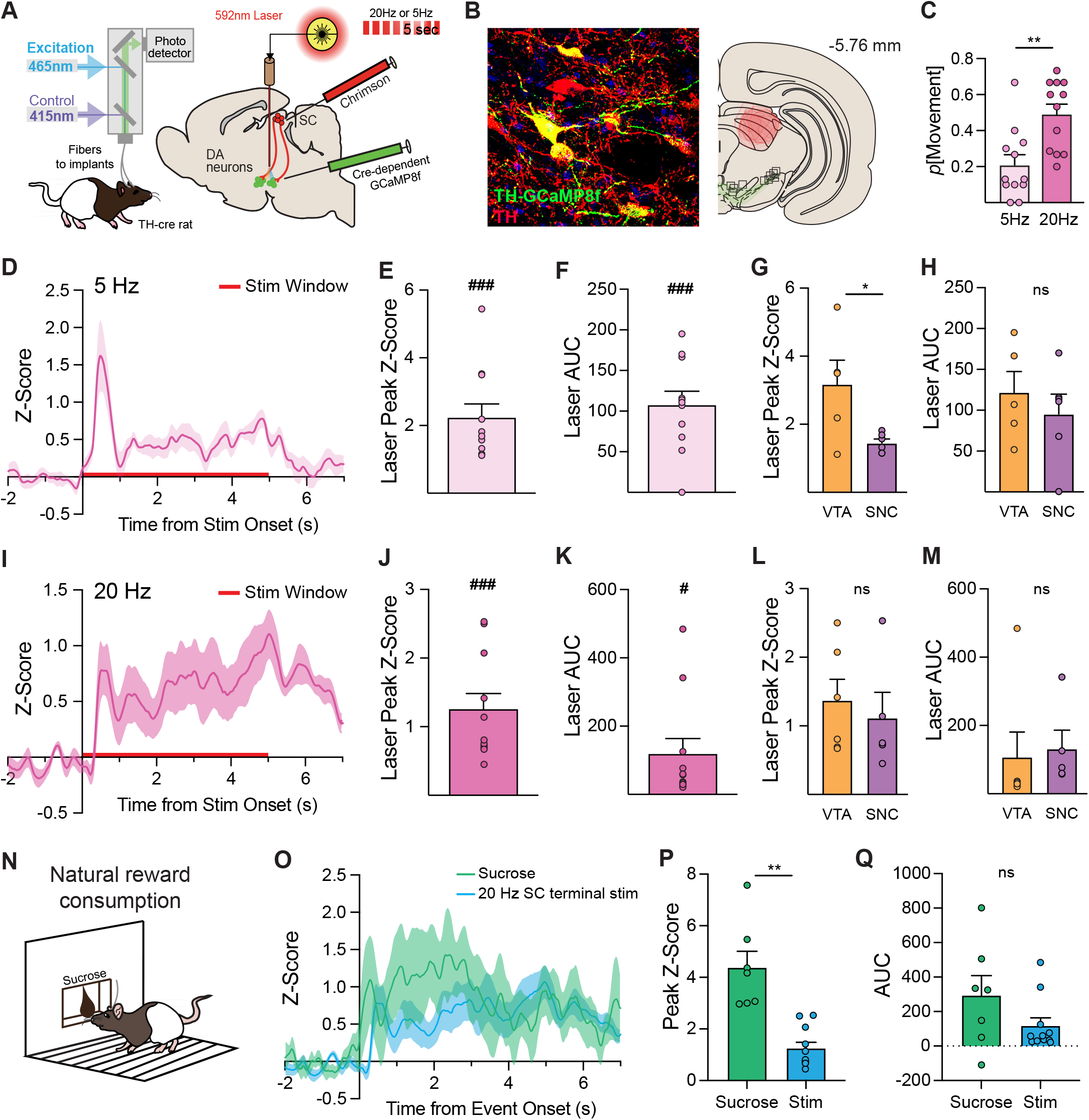
Superior colliculus projections activate VTA/SNc dopamine neurons *in vivo*. A) Approach to target SC terminals with the red-shifted excitatory opsin ChrimsonR and dopamine neurons with a cre-dependent GCaMP8-coding virus in TH-cre rats, for simultaneous optogenetic stimulation and photometry recordings in the VTA or SNc. B) Histology image showing targeting of GCaMP to TH-positive neurons, and fiber placements in the midbrain. C) SC terminal stimulation evoked locomotion with increased intensity at 20 versus 5 Hz laser delivery (N=12). D) Z-score average trace of DA neuron fluorescence for all rats time locked to 5-Hz stimulation onset (N=11). Robust phasic DA neuron activation was seen, as measured by signal E) peak and F) area under the curve (AUC) measures. VTA (N=5) and SNc (N=6) DA neuron activity differed in G) peak signal, but not H) AUC. I) Z-scored trace of DA neuron fluorescence time locked to 20-Hz stimulation (N=11), which produced robust sustained DA neuron activation, as measured by signal J) peak and K) AUC measures. VTA (N=6) and SNc (N=5) DA neuron activity at 20 Hz did not differ in L) peak signal or M) AUC. N) Rats (N=7, 4 VTA, 3 SNC) consumed sucrose reward from a port in the chamber. O) Both sucrose and 20-Hz SC terminal stimulation evoked strong dopamine neuron responses. P) Sucrose evoked a larger peak dopamine signal, Q) but a similar overall response based on AUC. **p<.01 (unpaired t test), *p<.05. ###p<.001 (one sample t test vs 0), ##p<.01 (vs 0), #p<.05 (vs 0). Error bars depict subjects mean +/-SEM.

To compare these artificially-evoked dopamine signals to those in response to natural reward, a subset of rats was given the opportunity to drink sucrose solution from a reward port in the chamber, while dopamine neuron activity was recorded (**Fig. 2N**). Sucrose consumption evoked reliable dopamine neuron activity, which was qualitatively similar to the pattern of activity as the 20-Hz terminal stimulation. The peak signal for sucrose was significantly greater than the peak from 20-Hz stimulation (**Fig 2P**, t test, t(7.6)=4.609, p=.002) while the AUC values were not statistically distinct (**Fig. 2Q**; t(7.97)=1.43, p=.1909.

### Superior colliculus projections excite VTA GABA neurons *in vivo*

SC terminals have been shown to also make contact with GABAergic neurons (Comoli et al., 2003; Solié et al., 2022; Z. Zhou et al., 2019), which make up approximately 30% of neurons in the VTA (Nair-Roberts et al., 2008). To determine how SC input affects the activity of these neurons *in vivo*, we combined fiber photometry recordings of GABA neurons with optogenetic activation of SC terminals in the VTA. To target GABA neurons, we used a promoter-based viral strategy (Scott et al., 2023; Wakabayashi et al., 2019; Stelzner et al., 2024). One virus delivered cre-recombinase with transfection driven by the GAD1 promoter, while a second, cre-dependent virus delivered GCaMP8. Through this method we recorded the impact of SC terminal stimulation on GABA neurons in the VTA (**Fig 3A,B**). Stimulation at 5 Hz (**Fig. 3C**) produced a robust increase in GABA neuron activity (**Fig 3D,E**; one sample t test peak t(7)=8.15, p<.0001; AUC t(7)=5.044, p=.0015). Stimulation of SC terminals at 20 Hz (**Fig 3F**) also evoked robust GABA neuron activity (**Fig 3G,H**; t test peak t(6)=4.202, p=.0057; AUC t(6)=4.977, p=.0025).

**Fig 3.**
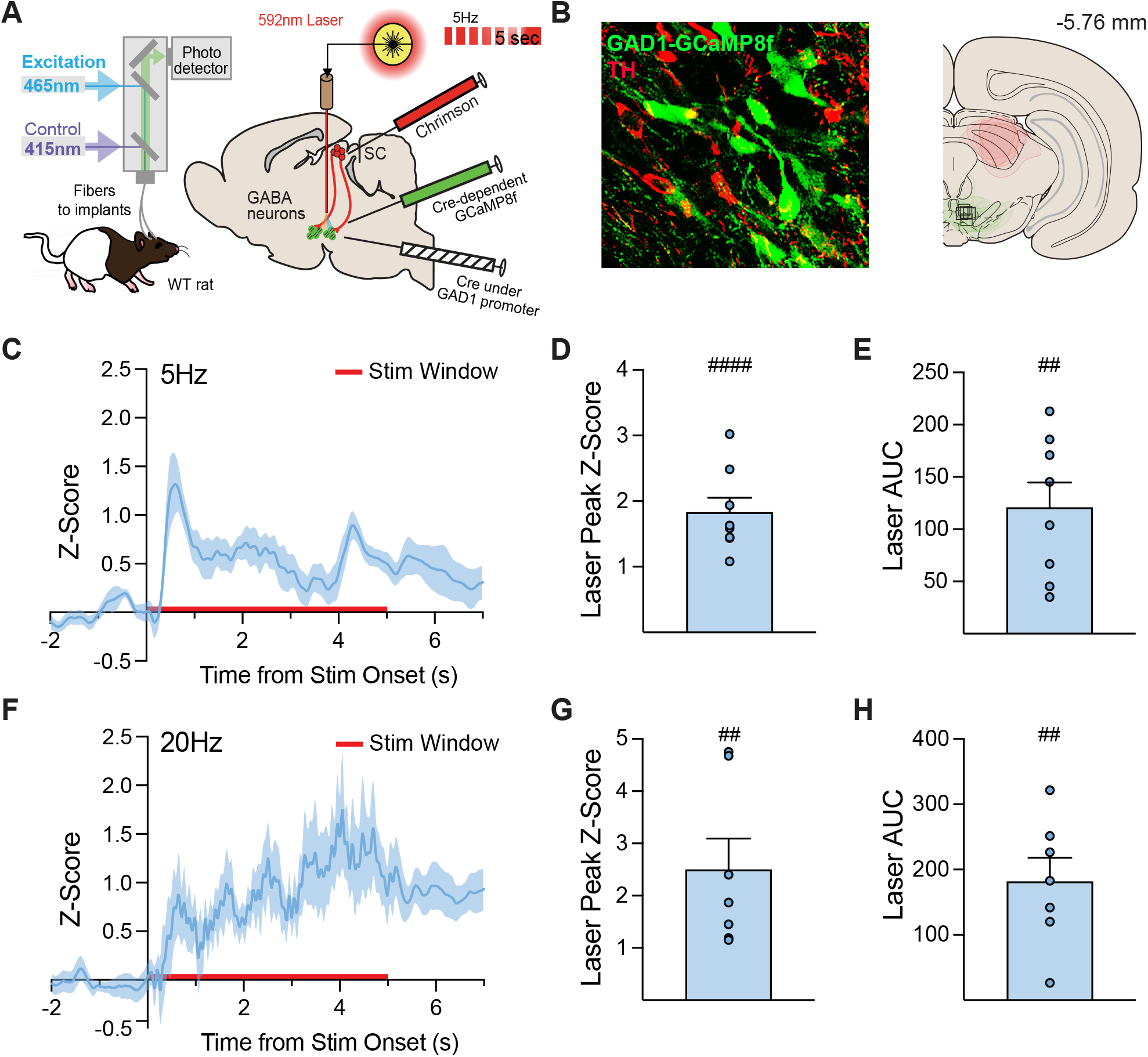
Superior colliculus projections activate VTA GABA neurons *in vivo*. Approach to target SC terminals with the red-shifted excitatory opsin ChrimsonR and GABA/GAD1+ neurons with a cre-dependent GCaMP8-coding virus in wild type rats, for simultaneous optogenetic stimulation and photometry recordings in the VTA. B) Histology image showing GAD-GCaMP targeting in TH negative neurons and fiber placements in the midbrain. C) Z-score averaged trace of GABA neuron fluorescence time locked to 5-Hz stimulation (N=8). Robust phasic GABA neuron activation was seen, as measured by signal D) peak and E) AUC measures. F) Z-scored trace of DA neuron fluorescence time locked to 20-Hz stimulation (N=7). Robust sustained GABA neuron activation was seen, as measured by signal G) peak and H) AUC measures. ##p<.01 (one sample t test vs 0), ####p<.0001 (vs 0). Error bars depict subject mean +/-SEM.

Taking our results (**Fig 2 and 3**) together, we show that SC neurons projecting to the ventral midbrain can evoke movement invigoration and excite both dopamine and GABA neurons in the VTA/SNc.

### Superior colliculus projections to VTA/SNc do not drive Pavlovian cue learning

Our previous studies (Engel et al., 2024; Saunders et al., 2018) have shown that brief stimulation of dopamine neurons in the SNc and VTA is sufficient to create conditioned responses to an associated, previously neutral cue. We hypothesized that SC projections to dopamine neurons contain sensory information about salient cues, and that these projections may therefore drive Pavlovian conditioned responses. To test this, we employed an optogenetic Pavlovian conditioning procedure. We expressed ChR2 in deep layer SC neurons and implanted a stimulating optic fiber over the ipsilateral VTA, SNc, or a control region elsewhere in the midbrain (**Fig 4A,B**). Rats were given 12 sessions wherein presentations of a neutral visual stimulus (cue light, 7 sec) were paired with brief optogenetic excitation of SC terminals (473nm laser, 5 sec, 20Hz). As with our previous studies using this approach (Saunders et al., 2018), no behavioral responses were required to obtain either cue or laser presentations, and no external rewards were given, in order to assess the inherent conditioning power of the circuit. Video recordings of conditioning sessions were first analyzed via hand scoring, with experimenters blind to the animal’s implant target. We focused on cue orientation/approach (**Fig 4D**), a common cue-evoked behavior seen for direct dopamine neuron stimulation, and in response to naturally conditioned stimuli.

**Fig 4.**
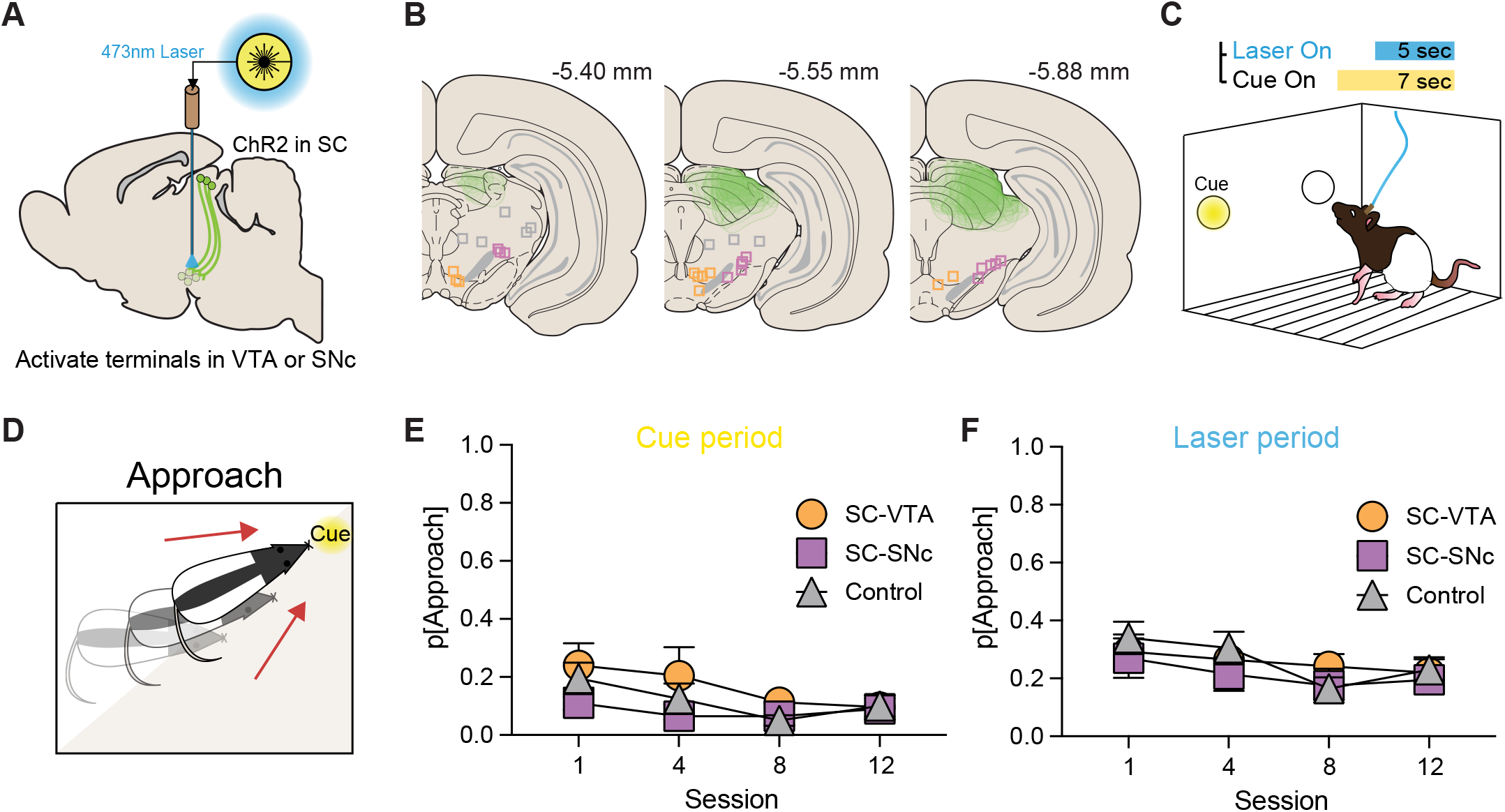
Activation of SC projections to the VTA/SNc does not drive Pavlovian cue conditioning. A) Approach for targeting of SC terminals in the ventral midbrain. ChR2-YFP expressed in deep layer SC neurons, and B) an optic fiber was implanted over the ipsilateral VTA (SC-VTA, n = 6), SNc (SC-SNc, n = 8), or a control region (Control, n = 8). C) Optogenetic Pavlovian conditioning paradigm, where a neutral cue was paired with optogenetic activation of SC terminals. D) Instances of cue approach behavior were measured via inspection of video recordings during conditioning. E) During the cue-only period, corresponding to the first 2-sec of cue presentations, minimal approach behavior occurred across all groups. F) During the laser period, corresponding to the final 5 sec of the cue when laser was also delivered, approach was similarly low for all groups. Error bars depict SEM.

Contrary to our prediction, SC terminal stimulation did not drive reliable cue-conditioned behavior. Across training, approach behavior in response to the cue alone remained at a low level for SC-VTA and SC-SNc stimulation groups, similar to the level seen for a control group where laser delivery was targeted outside of the ventral midbrain (**Fig 4E**; no effect of stim group, F(2,19)=2.01, p=.16). The approach behavior that did occur to the cue diminished across training (main effect of session, F(2.31,43.28)=3.796, p=.023). Focusing on the period of laser stimulation, we again did not see emergence of cue approach for either SC-VTA or SC-SNc groups (**Fig 4F**; no effect of group, F(2,19)=.516, p=.605). Thus, in stark contrast to general dopamine neuron activation (Engel et al., 2024; Saunders et al., 2018), activation of SC neurons projecting to dopamine-rich brain regions, while evoking dopamine neuron activity, does not imbue neutral stimuli with conditioned value to drive learning.

### Superior colliculus projections to VTA/SNc evoke head and body turning

Based on our above finding (**Fig 2**), where unsignalled SC terminal optogenetic activation in the VTA/SNc evoked movement, we comprehensively examined movement during our optogenetic Pavlovian conditioning. Despite not clearly driving cue learning, in these studies, we also found that SC terminal activation produced a consistent motor output, in the form of a head/upper body turning. This behavior, which we labeled as a “craning” response, was always directed contralateral to the stimulation hemisphere. For an initial analysis, we operationalized a crane occurring if the rat’s head turned at least 90 degrees, based on a quadrant axis overlaid on the video frame (**Fig 5A**). Critically, craning behavior did not reliably emerge during the cue-only period before the laser was turned on (Fig 5B; no effect of stim group, F(2,19)=.698, p=.51). Instead, we saw reliable head turning/craning only during the period of laser stimulation. This turning occurred strongly in the SC-VTA and SC-SNc subjects, relative to controls (**Fig 5C**; main effect of stim location, F(2,19)=24.05, p<.0001), which showed no consistent behavioral response of any kind. While SC-SNc rats exhibited substantial craning, the probability of craning was even higher in SC-VTA rats, an effect that was clearest by the end of the conditioning paradigm (post hoc comparison VTA vs SNc session 12, p=.0236). Although we saw a qualitative increase in the probability of craning in the VTA group across conditioning (**Fig 5C**, there was no overall change in craning likelihood among groups (no main effect of session, F(2.45,46.6)=1.66, p=.196).

**Fig 5.**
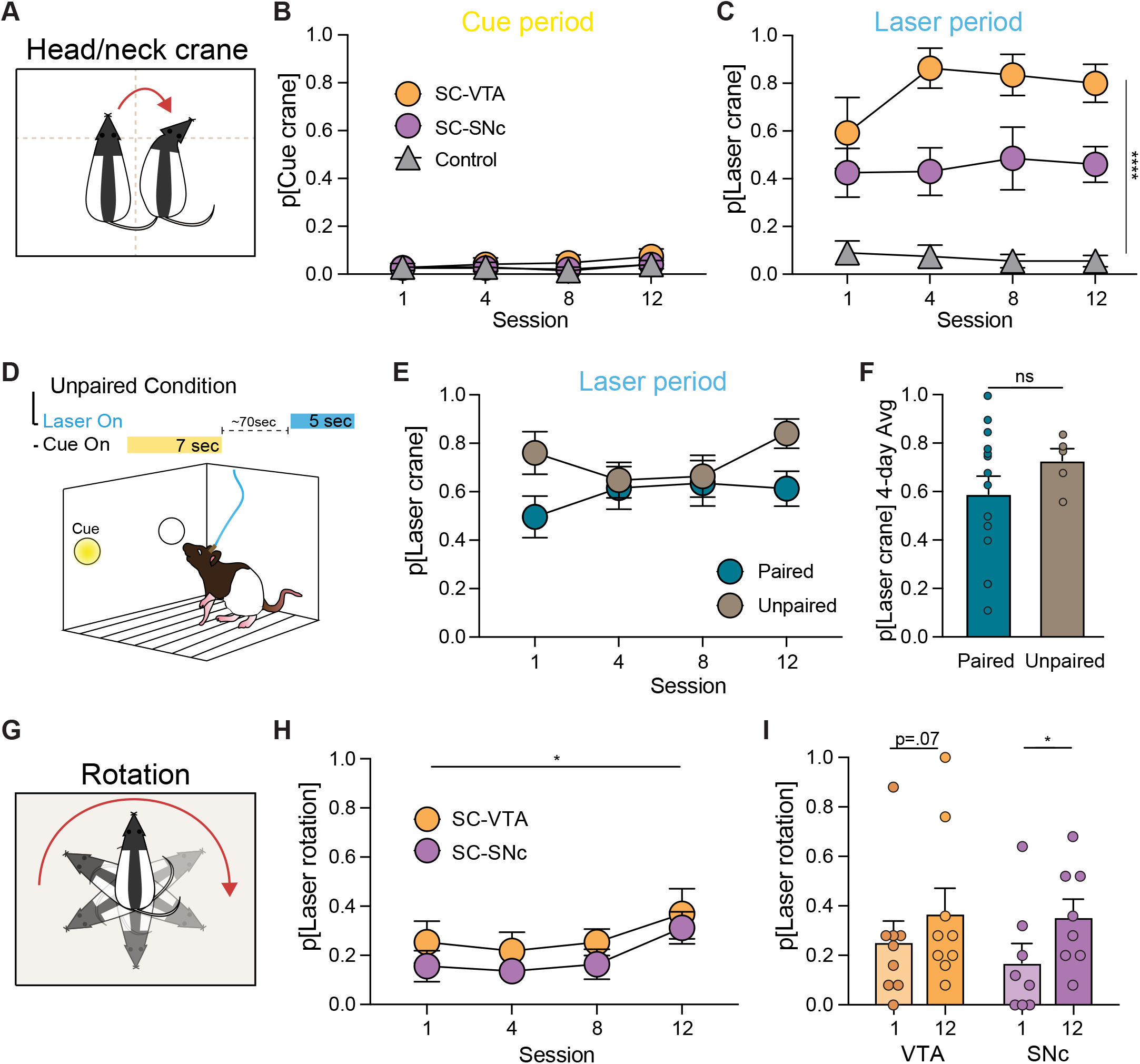
SC terminal stimulation in the VTA and SNc evokes head/neck turning independent of cue conditioning. A) Schematic illustrating craning behavior, which was defined as a head deflection of at least 90 degrees contralateral to the stimulation hemisphere. B) Craning behavior did not occur during the cue-only period (before laser onset). C) During the laser period, craning was robust in the SC-VTA (n=6) and SC-SNc (n=8) groups relative to controls (n=8). D) Unpaired optogenetic conditioning procedure, where cue and laser presentations were separated by a variable interval, for a separate cohort of rats (n = 5). E) Unpaired rats exhibited a similar probability of craning as paired subjects across training, and F) when data was collapsed across all analysis sessions. G) In addition to head craning, full body rotation contralateral to the stimulation hemisphere was quantified. (H,I) Rotations during the laser period occurred on a subset of trials, increasing in probability across conditioning sessions for SC-VTA and SC-SNc rats. ****p<.0001, *p<.05. Error bars depict SEM.

In our cue-laser paired rats, we saw that turning only clearly occurred during the laser portion of the cue presentation, and not to the cue alone. To examine the effect of the cue presentation on craning, we ran a separate cohort of rats (N=5, 3 SC-VTA, 2 SC-SNc) through an unpaired version of the optogenetic conditioning paradigm. In this version, the same number of cue and laser presentations were given across 12 sessions, but each cue and laser event were separated in time and never overlapped (**Fig 5D**). These unpaired rats also showed robust craning behavior to laser but not cue presentations. Across conditioning, there was no significant difference in the probability of craning for paired versus unpaired rats during laser stimulation (**Fig 5E**, no effect of condition, F(1,17)=1.14, p=.301). Averaged across sessions, unpaired craning was nominally higher than paired, but not significantly different (**Fig 5F**, unpaired t test, t(17)=1.07, p=.301). Thus, craning behavior was evoked by SC terminal stimulation, and was not a byproduct of cue-evoked learning.

Craning responses mostly reflected an upper body turn, but in some cases this behavior became a full body rotation, which we also quantified (**Fig 5G**). As with other behaviors, rotations occurred specifically during laser periods. Across all rats in the SC-VTA and SC-SNc groups, while rotations only occurred on a subset of trials, rotation probability increased across optogenetic conditioning (**Fig 5H**; main effect of session, F(2.05,34.8)=3.96, p=.027), suggesting an overall potentiation in the effect of SC terminal stimulation on turning behavior. Within the SC-VTA group, there was a trend for an increase in rotations from the first to last session (**Fig 5I**, paired t test, t(8)=2.01, p=.079), and a significant increase in the SC-SNc group (t(7)=3.135, p=.0165).

Above, we showed that SC neurons activate dopamine neurons *in vivo* (**Fig 2**). Increased dopamine activity is often associated with generalized locomotion (Howe & Dombeck, 2016), and repeated dopaminergic activation via pharmacologic (Magos, 1969), optogenetic (Lobo et al., 2010; Saunders et al., 2018), or chemogenetic (Ferguson et al., 2011) manipulation can sensitize locomotion. Therefore, we next examined locomotion more generally, irrespective of cue and laser presentations. To quantify the movement, we used the markerless pose estimation tool, DeepLabCut (Mathis et al., 2018) to identify rat body parts within the chamber. Using this positional data, we calculated the average speed of rats for each conditioning session (session length ∼45 min). This analysis revealed that SC terminal stimulation did not produce a general state of hyperactivity. Average movement speed was similar for SC-VTA, SC-SNc, and control rats (No effect of stim location, F(2,15)=.349, p=.717; avg speed: VTA 6.6 cm/s, SNC 6.26 cm/s, control 5.56 cm/s). Further, repeated stimulation did not produce locomotor sensitization, as average speed did not change across conditioning sessions (F(2.16,31.7)=1.30, p=.287).

### SC projections transiently shape postural reorientation

In **Fig 5**, we quantified the probability of craning based on an operational cutoff of >90 degrees turned. Across animals, and trial to trial, the extent of turning was variable, and turns were often below our cutoff threshold. To assess this in more detail, we used pose data from our DLC model to determine the degree of reorientation during cue and laser periods (**Fig 6**). A vector running from the middle of the back to the front of the animal was calculated. The vector’s relative position was compared frame to frame, and the cumulative angle of turning was quantified (**Fig 6B,C**). This approach revealed that upper body turning was tightly coupled to the laser delivery window in both SC-VTA (**Fig 6D**) and SC-SNc (**Fig 6E**) rats, with no turning occurring for anatomical control subjects (**Fig 6F**).

**Fig 6.**
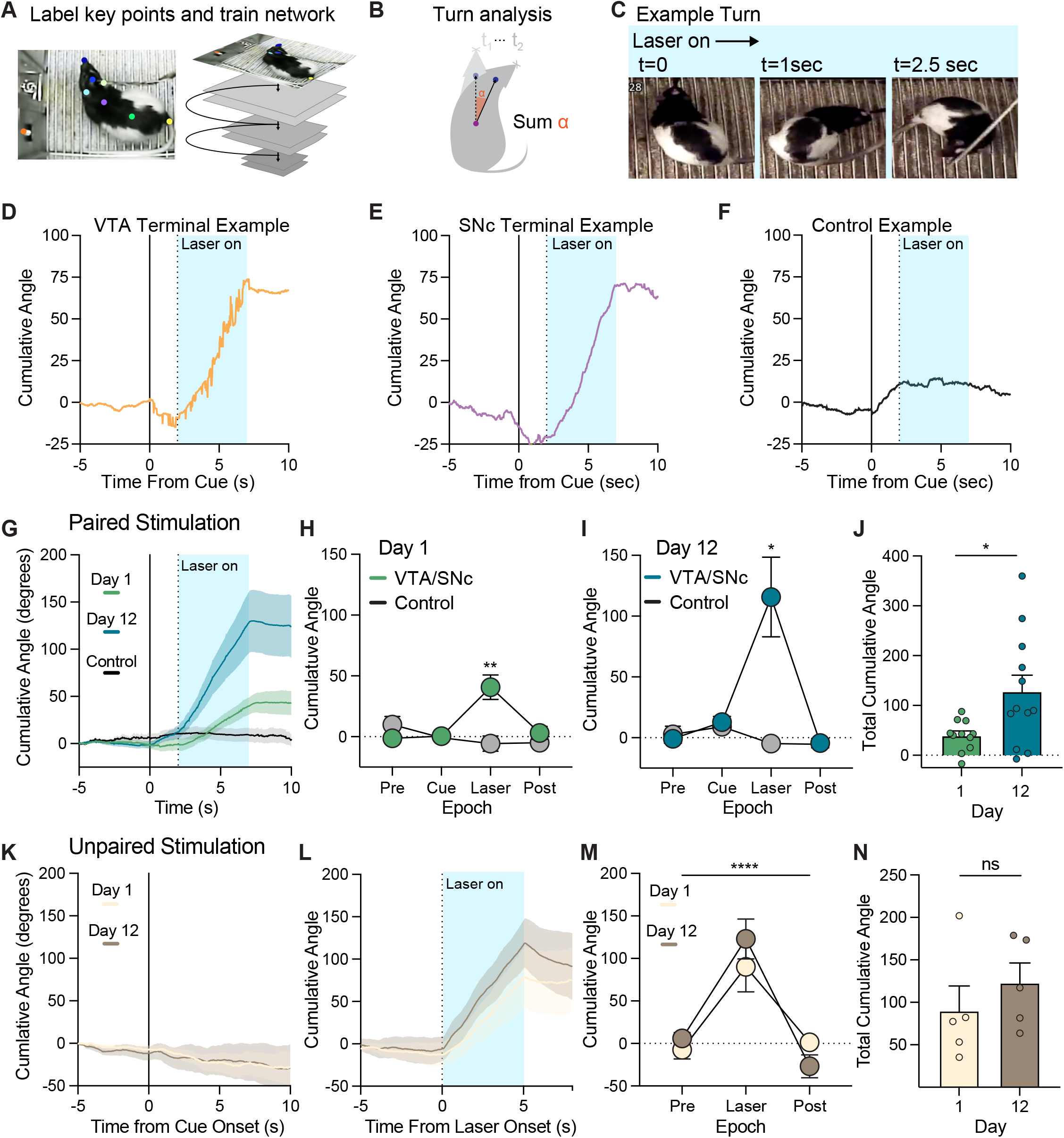
SC projections to VTA/SNc transiently shape head turning. A) Rats’ body parts and static elements of the chamber were labeled in video frames from behavior sessions to train a neural network and acquire pose data using the DeepLapCut pipeline. B) A directionality vector was interpolated and tracked across video frames for calculation of turning angle. C) Example turn, showing a rat at three sequential timepoints during a laser stimulation event. Turning examples for D) SC-VTA, E) SC-SNc, and F) control subjects. The blue shaded zone corresponds to the laser stimulation window. A positive slope corresponds to an increase in the cumulative angle of deflection of the head vector contralateral to the hemisphere of laser stimulation. G) Averaged turning traces for paired SC-VTA/SNc rats (n=12) on the first and last day of conditioning versus controls. On the H) first and I) last sessions, turning occurred only during the laser period in VTA/SNc rats, and not in controls. J) Laser-evoked turning intensity increased for paired rats from the first to last day of conditioning. K) Average cumulative angle traces for unpaired VTA/SNc rats (n=5) show no turning during unpaired cue presentations, but L) robust turning during the laser windows. M) As with paired rats, turning was specific to the laser epoch. N) Unpaired rat turning increased qualitatively, but was not significantly higher on the last day of conditioning. ****p<.01, *p<.05. Error bars depict SEM.

Overall, the pattern and intensity of turning was similar for SC-VTA and SC-SNc rats, and so we collapsed these groups for remaining analyses. We first compared cumulative turning in the cue-paired rats, plotting angle deflection from cue onset and during the laser period for the first and last conditioning sessions (**Fig 6G**). The laser-evoked nature of turning was evident in this group data, where a consistent increase in the slope of traces indicates a cumulative increase in the angle of head/neck deflection, specifically during the laser window. Splitting the cumulative angle into epochs showed that significant turning only occurred during the laser window, and not in response to the cue alone, or following termination of laser stimulation, for both the first (**Fig 6H**; epoch by group interaction, F(3,45)=7.48, p=.0004) and last (**Fig 6I**; F(3,48)=6.02, p=.0014) session of conditioning, relative to controls. Further, we found that for the paired rats, the intensity of turning significantly increased across conditioning (**Fig 6J**; paired t test, t(10)=2.76, p=.02), consistent with the increase in rotations shown in **Fig 5**.

We separately analyzed turning in the unpaired rats. For these animals, no consistent turning occurred in response to cue presentations (**Fig 6K**). Laser delivery evoked consistent turning, which stopped upon laser termination, similar to the paired rats (**Fig 6L,M**; main effect of epoch, F(1.41,11.27)=28.57, p<.0001). While turning intensity was nominally higher on the last day of conditioning for unpaired rats, this difference was not statistically different (**Fig 6N**; paired t test, t(4)=1.37, p=.243). However, when laser-evoked turning for paired and unpaired rats was averaged together, we found a strong significant increase in turning intensity from the first to last session (paired t test, t(15)=2.95, p=.0098). Collectively, these data indicate that activation of SC terminals in the VTA and SNc promotes a specific, focal behavioral pattern that includes reorientation of the head and neck.

### Superior colliculus projections to VTA/SNc do not drive reinforcement or valence assignment

Previous studies show that direct activation of VTA/SNc dopamine neurons supports robust reinforcement (Engel et al., 2024; Fraser et al., 2023; Ilango et al., 2014; Saunders et al., 2018; Tsai et al., 2009), inducing vigorous response patterns to obtain activation. Given our finding that SC neurons projecting to the VTA/SNc activate dopamine neurons, we next asked if stimulating these VTA/SNc inputs supports reinforcement. A subset of rats (SC-VTA = 4, SC-SNc = 4, Control = 2) were trained in an intracranial self stimulation (ICSS) task where active nose pokes resulted in delivery of a brief (1-sec) laser stimulation across three sessions (**Fig 7A**). Overall, rats failed to reliably engage in self-stimulation behavior (**Fig 7B**; no effect of nose poke, F(1,14)=.417, p = 0.5309). When examined separately, we found that SC-SNc rats self-stimulated nominally more than either SC-VTA or control subjects (**Fig 7C**; effect of implant location, F(2,14)=5.02, p = 0.023), but this reflected an increase in both active and inactive responses, and behavior was very low overall (∼10 nose pokes/session).

**Fig 7.**
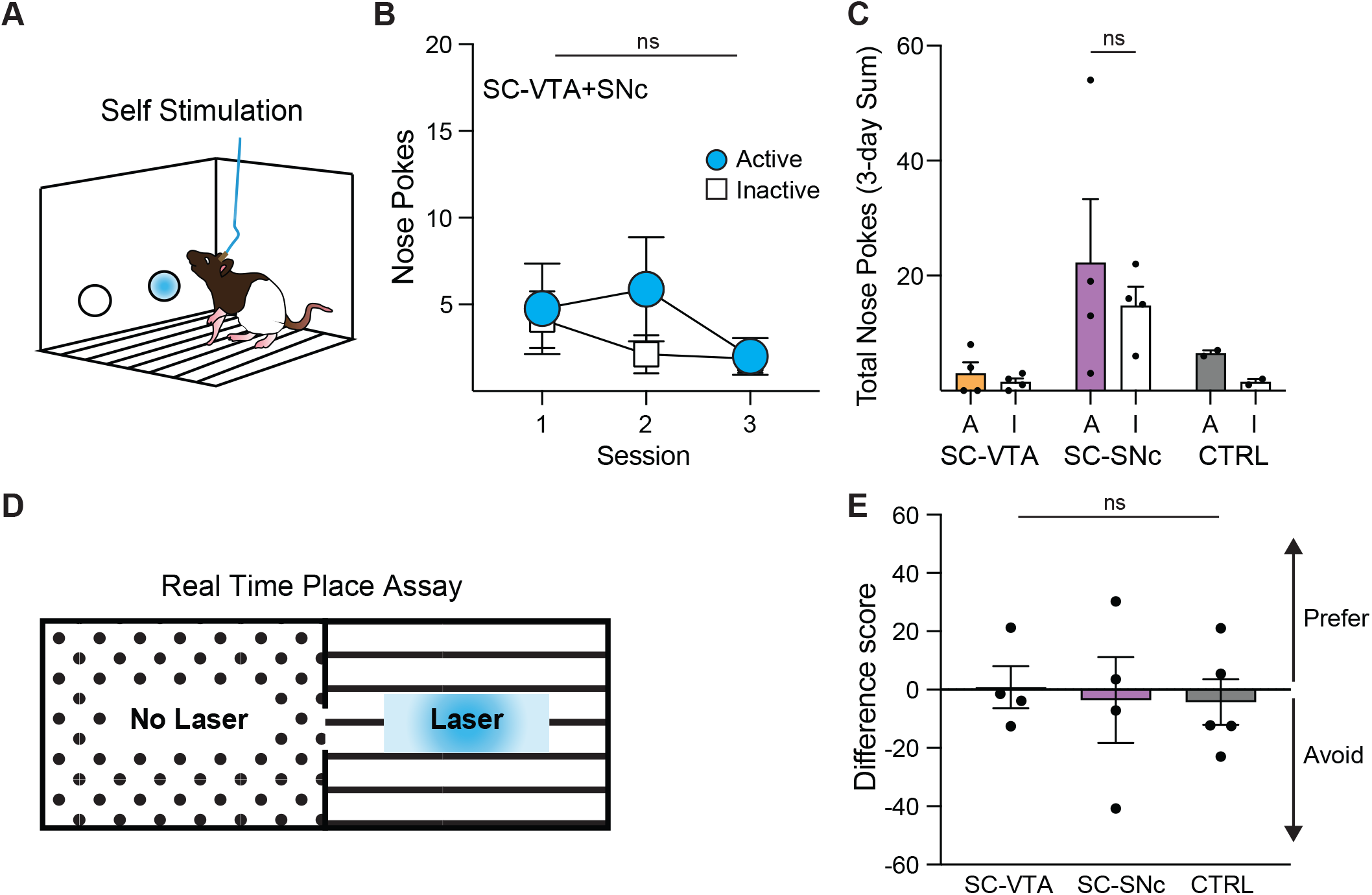
SC terminal stimulation does not support reinforcement or signal valence. A) Intracranial self stimulation (ICSS) set up. B) Nose behavior was low overall (n=8). C) Plotted separately, SC-VTA (n=4), SC-SNc (n=4), and control (n=2) groups exhibited low responding, failing to discriminate between active and inactive nose pokes. Active = A, Inactive = I. D) Real time place assay paradigm. E) No preference or avoidance was observed for any group. Error bars depict SEM.

Finally, we investigated if stimulation of SC terminals in the VTA/SNc drives an overall positive or negative valence, using an optogenetic real time place assay. Rats could move freely between two distinct chambers while tethered to a laser (**Fig 7D**). When >50% of the body crossed into the laser-paired chamber, SC terminals were optogenetically activated. Laser stimulation ceased when they crossed back into the neutral chamber. A difference score was calculated for each rat, based on the amount of time they spent in the stimulation chamber compared to the control chamber. A value of 100 would indicate the animal spent 100% of their time in the stimulation paired chamber, whereas a value of -100 would indicate the animal spent 100% of their time in the neutral chamber. There was no effect of stimulation location on difference score (**Fig 7E**; F(2,10)=.007, p = 0.9301). There was no side preference for any group, and all rats spent a relatively equal amount of time in both chambers, suggesting that stimulation was not strongly rewarding or aversive.

### Dopamine neurons receiving input from the SC promote turning but not reinforcement

Our results show that activation of SC terminals in the VTA/SNc invigorates turning behavior but does not drive learning and reinforcement. Given that SC inputs to the VTA/SNc target and activate both dopamine and GABA neurons, next we sought to isolate a possible dopamine-specific role in the turning phenotype. To do this, we injected an AAV1-flp virus into the SC of TH-cre rats, delivering flp recombinase transynaptically to the neurons that SC neurons innervate. We then injected a virus coding for ChR2-YFP under the control of both cre and flp recombinases into the ipsilateral VTA/SNc (**Fig 8A**). Thus, neurons containing both cre (dopamine neurons) and flp (neurons receiving SC input) completed the recombination logic and expressed ChR2-YFP. Based on our earlier results, we found no consistent differences in the impact of SC terminal stimulation in the VTA versus the SNc. Therefore, we chose to target the lateral VTA/SNc border for general expression throughout the ventral midbrain. This viral approach resulted in strong YFP expression in dopamine neurons in the VTA and SNc (**Fig 8B**). We found terminal expression of YFP downstream, throughout the striatum (**Fig 8C**). While YFP terminals were clearly present throughout the entire striatum, expression was visibly brighter and denser in the dorsal striatum, relative to ventral striatum (**Fig 8D**). These results extend our initial anatomical work (**Fig 1**), and are consistent with other studies (Solié et al., 2022; Watabe-Uchida et al., 2012) indicating that SC neurons target striatal-projecting neurons, including dopamine neurons.

**Fig 8.**
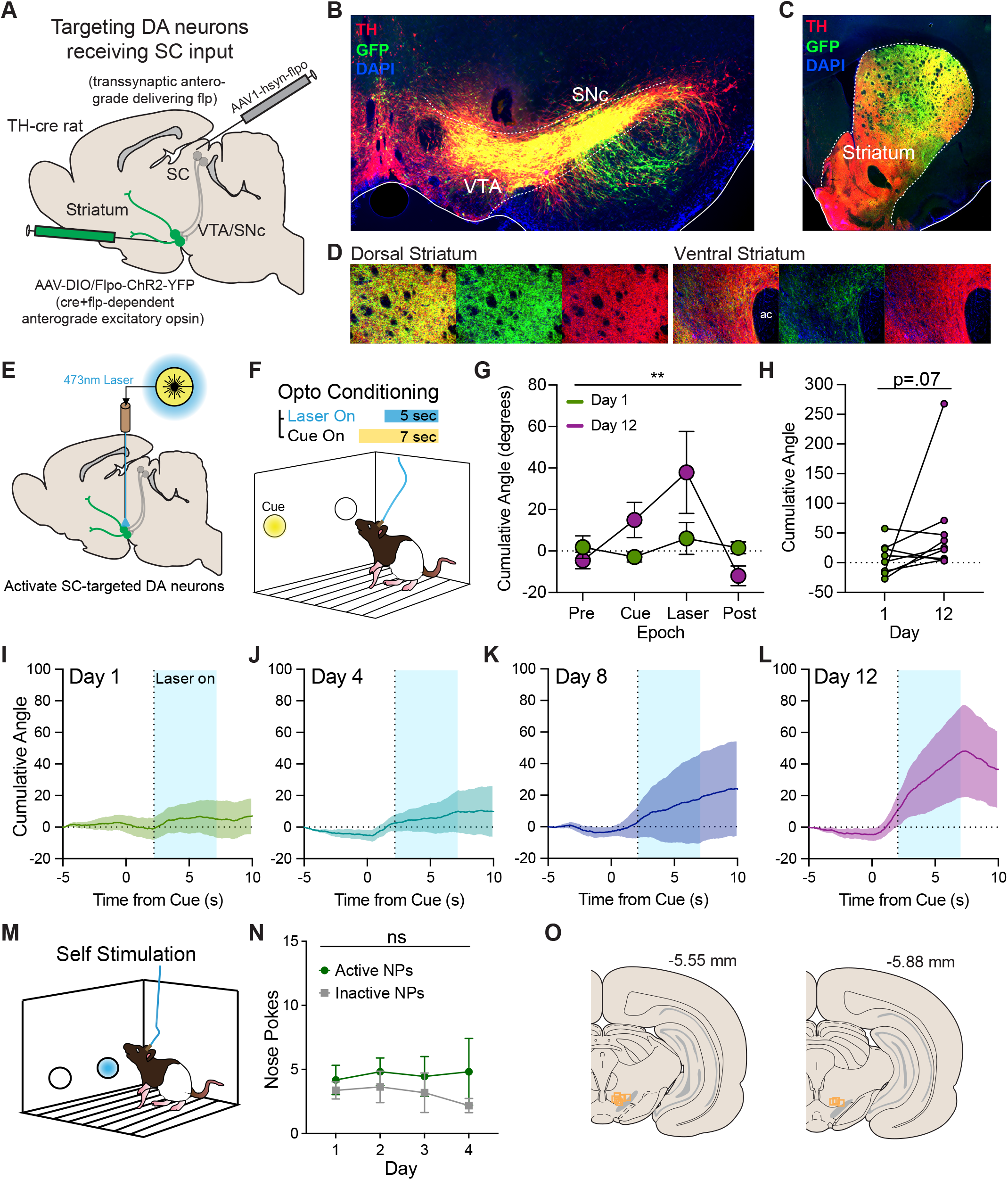
Dopamine neurons receiving input from the SC promote turning but not reinforcement. A) Viral approach for targeting dopamine neurons receiving direct input from the SC. In TH-cre rats, an AAV1-flp vector was injected into deep layer SC, followed by injection of a cre and flp-dependent virus coding for ChR2-YFP into the ipsilateral VTA/SNc. B) This resulted in strong expression of YFP in dopamine neurons, C) including those innervating the striatum. D) Zoomed in images showing YFP expression with TH counterstain in the dorsal and ventral striatum. E) Rats (N=9) with ChR2 expressed in dopamine neurons receiving SC input completed the F) optogenetic conditioning paradigm, where a cue predicted laser delivery. Turning behavior was quantified as the cumulative angle reached during cue/laser presentations. G) Averaged cumulative angle for trial epochs for the Day 1 versus 12 of training, where an increase during the laser period was evident. H) Total turning marginally increased from Day 1 to 12. I-L) Averaged cumulative angle traces across the trial period for Day 1, 4, 8, and 12, showing a steady increase in turning during laser. M) Rats were given the opportunity to nose poke for optogenetic stimulation of dopamine neurons that receive input from the SC. N) Across 4 training sessions, active and inactive nose pokes remained at a low level. O) Fiber optic placements. **p<.01. Error bars depict SEM.

Next, we conducted the optogenetic Pavlovian conditioning procedure, as described above, in rats (N=9) expressing ChR2 in dopamine neurons that are targeted by SC input (**Fig 8E,F,O**). Using pose data from our DLC model, we quantified turning behavior across 12 conditioning sessions. While minimal turning was seen on day 1, by the end of conditioning, there was a clear increase in cumulative angle during laser delivery (**Fig 8G**; main effect of epoch, F(3,24)=5.036, p=.0076). For most rats, total turning increased in magnitude from Day 1 to Day 12, with a trend for an overall average increase (**Fig 8H**; paired Wilcoxon test, p=.0742). We plotted the average cumulative angle trajectories relative to cue and laser onset for Days 1, 4, 8, and 12 (**Fig 8I-L**), which showed a general progression of greater turning across training. Next, rats were given the opportunity to nose poke to receive optogenetic activation of dopamine neurons receiving SC input, in an ICSS paradigm as above (**Fig 8M**). Across 4 sessions, rats failed to reliably engage in self-stimulation behavior, maintaining a low level of responding that did not discriminate between active and inactive nose pokes (**Fig 8N**; no effect of nose poke, F(1,10)=1.245, p=.291). Collectively, these studies show that SC neurons strongly innervate dopamine neurons and activation of this dopamine subpopulation is sufficient to induce turning, but not reinforcement.

## DISCUSSION

We investigated the anatomical organization and functional role of superior colliculus neurons projecting to the dopaminergic midbrain. Our primary findings are: 1) Deep layer SC neurons anatomically interface with dopamine neurons in the VTA and SNc. 2) SC projections excite dopamine neurons in the VTA and SNc, and GABA neurons in the VTA, *in vivo*. 3) Despite this, optogenetic excitation of SC projections to the VTA and SNc was not sufficient to drive Pavlovian learning about associated visual cues, nor did it promote reinforcement or aversion. 4) Instead, we found that stimulation of SC inputs to the VTA/SNc spurred head turning and body reorientation, and this motor effect potentiated with time. 5) Finally, we show that direct activation of dopamine neurons receiving synaptic input from the SC was sufficient to produce head turning, but not reinforcement. These results offer insights into a number of questions related to movement, learning, and attentional mechanisms impinging on ventral midbrain circuitry.

We find that deep layer SC neurons project to and excite dopamine neurons in the VTA and SNc. This is consistent with previous studies (Comoli et al., 2003; Dommett et al., 2005; May et al., 2009; McHaffie et al., 2006; Redgrave, Coizet, Comoli, Mchaffie, et al., 2010; Pradel et al., 2025), and we extend that work to show this process happens *in vivo* in behaving rats. We found some qualitative differences in the pattern and timing of evoked dopamine signals depending on the frequency of SC input. This suggests SC may exert nuanced control over the dopamine system, potentially, along with other inputs, modulating tonic and phasic-related activity modes in different behavioral contexts (Berke, 2018; Bromberg-Martin et al., 2010b; Grace et al., 2007; Tian et al., 2016; Wise & Robble, 2020; Pradel et al., 2025). Previous electron microscopy results show that SC projections to the ventral midbrain synapse on non TH-containing neurons (Comoli et al., 2003; Redgrave, Coizet, Comoli, Mchaffie, et al., 2010), including the notable population of GABA neurons that reside in the VTA (Nair-Roberts et al., 2008). We find that the SC directly activates VTA GABA neurons, in line with more recent studies using genetic tools (Z. Zhou et al., 2019; Pradel et al., 2025). The local interplay between GABA and dopamine neurons receiving input from the SC is unclear, but our results suggest that the bulk effect of SC-GABA excitation is not to simultaneously inhibit local dopamine neurons. One possibility is that SC inputs target GABA neurons that primarily project out of the VTA, including to frontal regions like the striatum or cortex (Solié et al., 2022; Taylor et al., 2014; W.-L. Zhou et al., 2022; Z. Zhou et al., 2019). Further, some SC neurons likely also interface with GABA neurons in the SNc and pars reticulata portion of the substantia nigra. It will be critical to further dissect the regional, microcircuit, and interhemispheric influences SC neurons have within the ventral midbrain.

The direct, general activation of dopamine neurons in the SNc and VTA drives strong learning, reinforcement, and value assignment (Coizet et al., 2007; Engel et al., 2024; Fraser et al., 2023; Huang et al., 2021; Ilango et al., 2014; Saunders et al., 2018; Tsai et al., 2009). Despite clearly activating dopamine neurons in the VTA and SNc *in vivo*, here we found that optogenetic activation of SC projections to these regions does not drive Pavlovian conditioning or promote reinforcement. It is possible that the negligible effect of SC inputs on learning creation may be due to the co-activation of GABA neurons, which have classically been viewed as opposing dopamine neuron function (Tan et al., 2012). More recent work, however, suggests that VTA GABA neurons have a more active role in appetitive behaviors than previously thought, and may engage unique local and distal circuit mechanisms, independent of dopamine neurons (Lefner and Moghaddam, 2025; Stelzner et al., 2024; W.-L. Zhou et al., 2022). Critically, when we targeted just dopamine neurons that receive input from the SC for optogenetic activation, we saw turning behavior, but did not see evidence of learning and reinforcement. We note that in our study of dopamine neuron activation, while turning was clearly evoked, the intensity of this effect was weaker than that of the general SC terminal activation studies. Based on this, we hypothesize that direct optogenetic excitation of GABA neurons that receive SC input also likely evokes movement invigoration. It will be critical to further unpack the relative contributions of SC circuits innervating GABA versus dopamine neurons, as well as glutamate neurons and other co-releasing subpopulations (Prévost et al., 2024). A large literature now indicates that subpopulations of dopamine neurons subserve unique learning and motivational functions (Collins & Saunders, 2020; Cragg et al., 1997; Ikemoto, 2007; Keiflin et al., 2019; Lammel et al., 2008; Tian et al., 2016; Ungless & Grace, 2012; Watabe-Uchida et al., 2012). One possibility is that SC may activate a subset of VTA/SNc neurons distinct from populations activated in previous studies demonstrating motivation and reward. Along these lines, our data provide further support for the notion that unique subpopulations of dopamine neurons may predominantly control movement versus reward (Engelhard et al., 2019; Howe & Dombeck, 2016; Greenstreet et al., 2025; Azcorra et al., 2023; Mantas et al., 2024). It is unclear if SC targeted dopamine neurons have specialized projection patterns, as we show that VTA/SNc neurons receiving SC input project broadly to striatum and ventral pallidum. Further investigation of this tectum-meso/nigro-frontal network is needed to support a more complete view of the likely multiple channels of influence collicular systems have on adaptive behavior (Chevalier & Deniau, 1984; Cover et al., 2024; Lak et al., 2020; Lee et al., 2020; Lee & Sabatini, 2021; Mana & Chevalier, 2001; Pradel et al., 2025), and disease (Bellone et al., 2024).

While many dopamine neurons are thought to encode reward prediction errors, individual circuit inputs do not transmit all features of reward-prediction errors independently. Instead, they are thought to signal mixed and distributed functions that differentially contribute features of learning (Beier et al., 2015; Bromberg-Martin et al., 2010a; Engelhard et al., 2019; Farassat et al., 2019; Lerner et al., 2015; Tian et al., 2016). This suggests that an integrative computation of the complete input activity to a dopamine neuron accompanies learning. Our results add to this notion, to show that a sensorimotor input from the SC is insufficient to drive learning on its own. Collectively, this underscores an important conclusion borne out by our data: activation of dopamine neurons is not part and parcel with “complete” learning (Millard et al., 2024; Fraser et al., 2023; Engel et al., 2024), and not all stimuli that activate dopamine neurons necessarily promote the same processes. The VTA and SNc are known to receive a variety of sensory and homeostatic afferents (Boughter et al., 2019; Watabe-Uchida et al., 2012), and it is therefore possible that for “complete” learning to occur, other components of a natural outcome (such as gustatory signals for food) may need to be incorporated into VTA/SNc to create a normal learned response. This is supported by our recording data, where sucrose consumption produced greater peak dopamine neuron activity compared to SC terminal stimulation. Indeed, it has been suggested that the neural correlates of learning, salience, movement, and value are likely somewhat distributed between inputs to dopamine centers (Bromberg-Martin et al., 2010b; Tian et al., 2016; Watabe-Uchida et al., 2012). It will be important to further explore what specific role SC inputs to dopamine neurons play within this set of computations. While cue-evoked dopamine signals are classically associated with reward-prediction error and reinforcement (Schultz, 1998), some have proposed that low-latency dopamine neuron activation acts as an alerting signal (Bromberg-Martin et al., 2010a; Redgrave et al., 1999; Schultz, 2007), conveying that an important stimulus is present and motor patterns should be engaged to explore it further. Our data are consistent with this notion, suggesting that SC-dependent dopamine activation in response to visual cues likely does not represent a reward prediction error or value signal (Amo, 2023; Berke, 2018; Lerner et al., 2021; Schultz, 1998).

SC activation of dopamine neurons in the VTA/SNc failed to create conditioned behavior to predictive cues, instead engaging specific movement patterns: rats turned during stimulation. This behavior showcases the control the SC has over head and neck (Cregg et al., 2020), and its general contributions to exploration. Reorientation of the head promotes survey of the environment, which is important for successfully identifying and navigating around rewarding or harmful stimuli. The deep layers of superior colliculus are known for coordinating exploratory behavior, via control of orientation (Isa et al., 2019; Suzuki et al., 2019; Villalobos & Basso, 2020) and attentional shifts (de Araujo et al., 2015; Goldberg & Wurtz, 1972; Hoy et al., 2019; Ignashchenkova et al., 2004; Krauzlis et al., 2013; White & Munoz, 2011; Yoshida et al., 2012). Our results, and others (Solié et al., 2022), indicate that this function is partially engaged via interface with the ventral midbrain. It will be important to further investigate the timing of *in vivo* control between SC and dopamine neurons. Here, our activation of SC inputs is presumably longer than what are often brief phasic signals elicited by sensory stimuli *in vivo* (Evans et al., 2018; K. Lee et al., 2020; Beltramo and Scanziani, 2019; Wang et al., 2020). As such, these inputs may normally function to induce rapid micro-adjustments in head position in all directions (Wilson et al., 2018), rather than body turning *per se*. The ability to orient to potentially salient stimuli at rapid latencies is an important step in learning about the environment and could provide insight into the function of SC-induced dopamine excitation. Notably, we saw evidence of a potentiation of the motor effect of SC terminal stimulation, and for the effect of stimulation of dopamine neurons receiving SC input. This suggests that some level of plasticity is engaged when SC inputs to the VTA/SNc are repeatedly active. Future studies will be useful for determining the specific conditions under which this occurs. Notably, VTA dopamine and GABA neurons are known to encode features of head movement and position, including the exertion of force to pitch the head (Engelhard et al., 2019; Hughes et al., 2020; Jiang et al., 2025). Our results suggest that this function may be at least partially governed by inputs from the SC. It will be important to investigate if that property is driven by SC inputs that encode head space (Wilson et al., 2018).

Strengthening head turning to reorient gaze to a particularly salient stimulus more rapidly over time could be one way that SC engages motor patterns (Basso et al., 2021; Hopp & Fuchs, 2004), via integration with the ventral midbrain, to promote attentional shifts across heterogeneous sensory learning contexts. The rapid coordination of movements in both investigatory and avoidance contexts is shown to be influenced by SC projections in important recent studies. For example, deep layer SC projections to the VTA/SNc contribute to fear-induced escape and avoidance (Sahibzada et al., 1986; Z. Zhou et al., 2019) while also signaling appetitive behaviors such as head orientation to conspecifics (Solié et al., 2022) and motion towards prey (Huang et al., 2021). When such inherently valuable stimuli are absent from the environment, as in our studies, we find that SC inputs drive dopamine neuron activity but do not artificially instantiate valuation. The ability of this pathway to create motor outputs to both positively and negatively valenced events is also consistent with the lack of either ICSS or place preference observed (Solié et al., 2022).

In conclusion, our results highlight a brain circuit that is important for guiding movement to redirect attention, via interaction with classic learning systems. We find that deep layer SC neurons rapidly excite dopamine neurons in both the VTA and SNc, and this activation spurs exploratory movement of space. While this circuit alone does not evoke associative learning, it may be important for priming key learning centers, like dopamine neurons, to important stimuli, for facilitation of association formation.

## METHODS

### Subjects

Male and female wild type and transgenic TH-cre (Witten et al., 2011) Long-Evans rats (N=60, 34 males, 26 females) were used in these studies. After surgery, rats were dual-housed with littermates and provided *ad libitum* access to food and water on a 0800 to 2000 light/dark cycle (lights on at 0800). All rats weighed >200g at the time of surgery and were between 3-6 months old at the time of experimentation. Experimental procedures were approved by the Institutional Animal Care and Use Committee at the University of Minnesota, Twin Cities and were carried out in accordance with the guidelines on animal care and use of the US National Institutes of Health.

### Viral vectors

To visualize superior colliculus (SC) terminals in the VTA and SNc, we used a GFP-expressing virus (AAV5-hsyn-GFP, Addgene #50465, 7×10^12^ vg/mL). For the visualization of synaptic contact between SC terminals and neurons in the VTA/SNc, a virus coding for membrane bound GFP and mCherry conjugated to the synaptic protein Synaptophysin was used (AAV5-hsyn-mGFP-2A-synaptophysin-mRuby, Addgene #71760, 7×10^12^ vg/mL). To visualize VTA/SNc neurons receiving direct SC input, we used an anterograde tracing approach exploiting the transsynaptic properties of AAV1 (Zingg et al., 2017). A cre-delivery virus (AAV1-hsyn-cre-WPRE, Addgene #105553, 1×10^13^ vg/mL) was injected into deep layer SC. Then, a cre-dependent virus expressing Tdtomato (AAV5-hsyn-DIO-tomato, Addgene #28306, 7×10^12^ vg/mL) was injected into the VTA/SNc, allowing for visualization of neurons receiving SC innervation. For combined optogenetic stimulation and photometry experiments, excitation of superior colliculus terminals was achieved via expression of the red-shifted excitatory opsin, ChrimsonR (Klapoetke et al., 2014; AAV5-syn-ChrimsonR-tdT; Addgene #59171, 7×10^12^ vg/mL), injected into SC. For photometry recordings of dopamine neurons, we injected a cre-dependent calcium indicator GCaMP8f (Zhang et al., 2023; AAV5-syn-DIO-jGCaMP8f-WPRE, Addgene #162379, 7×10^12^ vg/mL) into the VTA and SNc of TH-cre rats. For photometry recordings of GABA neurons, expression of GCaMP8f in GAD+ neurons was achieved through the combination of a cre-delivery virus GAD1-cre (AAV5-GAD1-cre, University of Minnesota Viral Vector and Cloning Core, 3×10^12^ vg/mL) and the cre-dependent GCaMP8f virus injected into the VTA of wild type rats, similar to previous studies (Scott et al., 2023; Wakabayashi et al., 2019; Stelzner et al., 2024). For general optogenetic conditioning experiments, channelrhodopsin (ChR2) was injected into the SC, for expression in terminals in the ventral midbrain (AAV5-hsyn-hChR2-EYFP, Addgene #26973 7×10^12^ vg/mL). To target dopamine neurons receiving synaptic input from the SC for optogenetic manipulation, a virus delivering flp-recombinase in an anterograde transynaptic fashion (AAV1-EF1a-flp, Addgene #55637, 7×10^12^ vg/mL) was injected into the SC in TH-cre rats. A second virus, coding for ChR2 expressed in neurons containing both cre and flp (Fenno et al., 2014; AAV5-Con/Fon-ChR2-YFP, Addgene #55645, 7×10^12^ vg/mL) was injected into the VTA/SNc.

### Surgical procedures

Viral infusions and optic fiber implants were carried out as previously described (Engel et al., 2024; Steinberg et al., 2014). Rats were anesthetized with 5% isoflurane and placed in a stereotaxic frame. Rats were administered saline, carprofen anesthetic, and cefazolin antibiotic subcutaneously. The top of the skull was exposed and holes were drilled for viral infusion needles, optic fiber implants, and four skull screws. Following skull exposure, surgery anesthesia was maintained at 1–3%. Viral injections were made using a microsyringe pump at a rate of 0.1 μl/min. Injectors were left in place for 5 min, then raised 100 μm dorsal to the injection site, left in place for another 10 min, then removed slowly. Implants were secured to the skull with dental acrylic applied around skull screws and the base of the ferrule(s) containing the optic fiber. At the end of all surgeries, topical anesthetic and antibiotic ointment were applied to the surgical site, rats were removed to a heating pad and monitored until they were ambulatory. Rats were monitored daily for 1 week following surgery. Optogenetic manipulations commenced 6–8 weeks after surgery.

For photometry recording of dopamine neurons, DIO-GCaMP8f was infused (0.5 uL at each target site for a total of 1 uL per rat) at the following coordinates from Bregma: VTA: posterior -5.4mm, lateral +0.8, ventral -8.1; SNc: posterior -5.4mm, lateral -2.6, ventral -7.3. To record GABA neurons, GAD1-cre and DIO-GCaMP8f were co-infused (0.5uL each, for a total volume of 1uL per rat) into the VTA (posterior -5.4, lateral +0.8, ventral -8.1). For optogenetic targeting of SC projections to the VTA and SNc, ChrimsonR (0.5 - 1uL per side, for a total volume of 1-2uL per rat) was also injected into deep layer SC at posterior -6.3, lateral +2.0, ventral -4.8. For simultaneous optogenetic activation of SC terminals and photometry recording of dopamine or GABA neurons, low-auto-fluorescence optic fiber implants (400-um glass diameter, Doric) were inserted just above viral injection sites at the following coordinates. VTA: posterior -5.4, lateral +1.0, ventral -7.8. SNc: posterior -5.4, lateral -2.6, ventral -7.2.

For SC terminal optogenetic Pavlovian conditioning, ICSS, and real time place assay experiments, ChR2 was delivered unilaterally (1uL total volume per rat) to deep layer SC (posterior -6.3, lateral +2.0, ventral -4.8). Optic fibers (300-nm diameter, custom made) were placed over ipsilateral VTA (posterior -5.4, lateral +0.8, ventral -7.5) or ipsilateral SNc (posterior -5.4, lateral -2.4, ventral -7.2). Subjects with targeting that ended up outside of the VTA/SNc were analyzed separately as anatomical comparisons.

For SC-targeted dopamine neuron optogenetic Pavlovian conditioning and ICSS experiments, AAV1-flp was delivered unilaterally (400nL) into deep layer SC. A cre and flp-dependent ChR2 coding virus was injected into the ipsilateral VTA/SNc (posterior -5.4, lateral +1.0, ventral -7.5).

### Optogenetic Stimulation

ChrimsonR studies used 590-nm lasers and ChR2 studies used 473-nm lasers (Dragon Lasers), adjusted to produce ∼2mW/mm^2^ light output from the tip of the intracranial fiber during individual 5-ms light pulses used in experiments. Light power was measured before and after every behavioral session to ensure that all equipment was functioning properly. For all optogenetic studies, optic tethers connecting rats to the rotary joint were sheathed in a lightweight armored jacket to prevent cable breakage and block visible light transmission.

#### Habituation and optogenetic Pavlovian training

Optogenetic Pavlovian training was based on the behavioral protocol described in previous studies (Engel et al., 2024; Saunders et al., 2018). Briefly, rats were first acclimated to the behavioral chambers (Med Associates), conditioning cue, and optic cable tethering in a ∼30-min session. During this session, 25 cue presentations, with no other consequences, were delivered on a 90-s average variable time (VT) schedule. In each of 12 subsequent conditioning sessions, rats in paired groups (N=22) were presented with 25 cue (light, 7s) – laser stimulation (100 5-ms pulses at 20 Hz; laser train initiated 2 s after cue onset) pairings delivered on a 200-s VT schedule. Rats in the unpaired group (N=5) also received 25 cue presentations and 25 laser trains per session, but an average 70-s VT schedule separated these events in time. These cues were never associated with another external stimulus (for example, food or water). The duration of laser stimulation was chosen to mimic the multi-second dopamine neuron activation observed *in vivo* when these subjects consumed natural reward, such as sucrose (Saunders et al., 2018). In all groups, cue and laser delivery were never contingent on an animal’s behavior and all rats received the same number of cue and laser events. Rats in the SC-targeted dopamine neuron experiment (N=9) received the same conditioning preparation.

#### Intracranial self stimulation (3-4, 1-h sessions)

A subset of rats from the SC terminal optogenetic conditioning experiment (N=12) were tethered to patch cables in the Med Associates boxes for self stimulation assessment. During these sessions, nose poke ports were positioned on the wall opposite of the cue lights and levers from previous phases. During 3 1-h sessions, pokes in the active port resulted in a 1-s laser train (20 Hz, 20 5-ms pulses, fixed-ratio 1 schedule with a 1-s timeout during each train), but no other external cue events, to assess the reinforcing value of stimulation itself. Inactive nose pokes were recorded, but had no consequences. Rats in the SC-targeted dopamine neuron experiment (N=9) underwent ICSS with the same conditions for 4 1-hr sessions.

#### Real time place assay (2, 30 min sessions)

A subset of rats (unique from ICSS, N=12) were tethered to patch cables and then habituated to a large open top arena (total arena size: 24”x16”x16”) split into two equally sized sections (12”x16”x16”). Each section had a uniquely textured floor and side panels (stripes or dots) distinguishing the two sections. Rats could move freely between both sections. Rats were placed in the center of the arena, and total time spent in each section was recorded throughout 30 min. Time was recorded as being in one chamber if > half of the rat’s body was located in the chamber. Times were totaled for each animal, and laser delivery was assigned to each rat’s less preferred chamber. The following day, rats were once again placed in the center of the chamber, and an experimenter observed the animals through a camera mounted above the arena for 30 min. When an animal crossed to the laser-paired side, stimulation was delivered (5mW, 5-ms pulses at 20 Hz) for the duration the rat was in said laser-paired chamber. When the rat crossed back to the non-laser side, stimulation ceased. To total the amount of time spent in each chamber, an experimenter blinded to the conditions watched the recorded videos and totaled time spent in each section. A difference score was calculated by subtracting the amount of time spent in the non-laser chamber from the amount of time spent in the laser-paired chamber.

### Video Scoring

Behavior during Pavlovian conditioning sessions was video recorded using cameras (Vanxse CCTV 960H 1000TVL HD Mini Spy Security Camera 2.8-12mm Varifocal Lens Indoor Surveillance Camera) positioned a standardized distance above each chamber. Videos from sessions 1, 4, 8, and 12 were scored offline by observers who were blind to the identity and anatomical target group of the rats. Each cue (7 s, 25 per session) and laser (5 s, 25 per session) event was scored for the occurrence and onset latency of the following behaviors. Locomotion was defined as the rat moving all four feet in a forward direction (that is, not simply lifting feet in place). Cue approach was defined as the rat’s nose coming within 1 in of the cue light (trials in which the rat’s nose was in front of the light when it was presented were not counted in the approach measure). Approach often involved the rat moving from another area of the chamber to come in physical contact with the cue light while it was illuminated. Rearing was defined as the rat lifting its head and front feet off the chamber floor, either onto the side of the chamber, or into the air. Rotation was defined as the rat making a complete 360-degree turn in one direction. Head turning/craning was defined as the rat’s head turning at least 90-degrees in one direction, based on visual assessment of the rat’s head/neck position relative to a grid overlaid onto the video frame.

### DeepLabCut-based behavioral analysis

Markerless tracking of rat body parts and position was conducted using the DeepLabCut (DLC) Toolbox (Mathis et al., 2018) and analysis of movement features based on these tracked coordinates was conducted in Matlab R2021b (Mathworks). DLC analysis was conducted on a Dell G7-7590 laptop running Windows 10 with an Intel Core i7-9750H CPU, 2.60Ghz, 16 GB RAM, and an NVIDIA GeForce RTX 2080 Max-Q 8GB GPU. DeepLabCut 2.1.10 was installed in an Anaconda environment with Python 3.7.7 and Tensorflow 1.13.1. Videos (944 x 480 resolution) were recorded with a sampling frequency of 30 frames per second using a Tiger Security Super HD 1080P 16-Channel DVR system.

### DeepLabCut model

For animal tracking we refined a network generated in our previous studies (Engel et al., 2024). Body parts and environmental features were labeled in 2090 frames from 35 videos (from 32 different animals) and refined the network by adjusting 807 outlier frames. 95% of these labeled frames were used for training. We used a ResNet-50 based neural network model for 1,030,000 training iterations. After final refinement and using a p-cutoff of 0.85, training error was 2.99 pixels and test error was 3.68 pixels. The body parts labeled included the nose, eyes, ears, fiber optic implant, midpoint of the shoulders, tail base, and an additional three points along the spine. Features of the environment were also labeled, including the 4 corners of the apparatus floor, two nose ports, two cue lights, two magazine ports, and 3 LED indicator lights when active.

To time-lock behavior to ongoing experimental variables (such as the onset of lights and laser stimulation) a separate network was trained to identify the onset of the cue light and the laser indicator light using 480 frames from 12 videos (10 different animals) and a 95% training dataset. We used a ResNet-50 based neural network model for 1,000,000 training iterations and the training error was 1.39 pixels and the test error was 1.56 pixels after final refinement of outlier frames.

DLC coordinates and confidence values for each body part and environment feature for every frame were imported to Matlab and filtered to exclude body parts/features from any frame where the confidence was < 0.7. To convert pixel distances to the real chamber dimensions, for each video, a pixel to cm conversion rate was determined. The distance (in pixels) between each edge of the environment floor and the diagonal measurements from corner to corner was measured, and these values were divided by the actual distance in cm. The mean of these values was then used as the conversion factor. Movement speed was calculated from the implant coordinates frame by frame using the formula: [distance moved (pix per cm) * framerate] to give movement speed in cm/s. For detailed analysis of postural turning, cumulative angle was calculated by adding the change in angle (degrees) between the vector between the shoulder point and the mid back point on the current frame and this same vector in the previous frame using the formula: angle= atan2(norm(cross(a,b)), dot(a,b)). Resulting angle values were normalized for the implant hemisphere so that increases in cumulative angle reflected the movement direction (contralateral to the implant).

### Fiber photometry

Dopamine neurons in the VTA and SNc of TH-cre rats (n=12) were transfected with a cre-dependent GCaMP8f and SC neurons were transfected via the red-shifted opsin ChrimsonR. VTA GABA neurons were targeted with GCaMP8f in a separate cohort of rats (n = 8). Implantation of optic fibers in the VTA and SNc allowed for simultaneous optogenetic activation of SC terminals and measurement of activity-dependent fluorescence in the same region (Kim et al., 2016; Saunders et al., 2018).

Photometry was conducted similar to previous studies (Engel et al., 2024; Lerner et al., 2015; Saunders et al., 2018). A fluorescence mini-cube (Doric Lenses) transmitted light streams from a 465-nm LED sinusoidally modulated at 211 Hz, and a 415nm LED modulated at 531 Hz. LED power was set at ∼50 μW. The mini-cube also transmitted light from a 590-nm laser, for optogenetic activation of ChrimsonR through the same low autofluorescence fiber cable (400 nm, 0.48 NA), which was connected to the optic fiber implant on the rat. GCaMP8f fluorescence from neurons below the fiber tip in the brain was transmitted via this same cable back to the mini-cube, where it was passed through a GFP emission filter, amplified, and focused onto a high sensitivity photoreceiver (Newport, Model 2151). Demodulation of the brightness produced by the 465-nm excitation, which stimulates calcium dependent GCaMP8f fluorescence, versus isosbestic 415-nm excitation, which stimulates GCaMP8f in a calcium-independent manner, allowed for correction for bleaching and movement artifacts. A real-time signal processor (RZP, TuckerDavis Technologies) running Synapse software modulated the output of each LED and recorded photometry signals, which were sampled from the photodetector at 6.1 kHz. The signals generated by the two LEDs were demodulated and decimated to 382 Hz for recording to disk. For analysis, both signals were then downsampled to 40 Hz, low-pass filtered at 6Hz, and a least-squares linear fit was applied to the 415-nm signal, to align it to the 465-nm signal. This fitted 415-nm signal was used to normalize the 465-nm signal, where ΔF/F = (465-nm signal – fitted 415-nm signal)/(fitted 415-nm signal). Laser presentation was time stamped in the photometry data file via a signal from the Med-PC behavioral program, and behavior was video recorded as described above. Normalized signals for each stimulation trial were extracted in a window from 5 s preceding to 10 s after laser onset, and Z-scored to the 5 s pre-laser period. Signal peak and area under the curve (AUC) values were calculated from the Z-scored traces by numerical integration via the trapezoidal method using the trapz function (Matlab).

For recording experiments, rats first underwent a day of tether training, wherein they were attached to optic cables and free to explore the chamber. Rats then received 2 counterbalanced stimulation-recording sessions, one with stimulation delivered at 20 Hz, and one with stimulation delivered at 5 Hz. In each of these sessions, 10 unsignaled laser trains (∼2mW 50 ms pulses, 5 sec train) were delivered on a 400-s variable time interval.

### Data collection, statistics, and analysis

Rats were randomly assigned to conditioning groups (paired, unpaired) following surgery. Behavioral data from optogenetic conditioning experiments was automatically recorded with Med-PC software (Med Associates) and analyzed using Prism 10.0. Video of conditioning sessions was recorded using a Tiger Security Recording System, and behavior data was analyzed in MATLAB. For manual video scoring, experimenters were blind to the anatomical group identity. Two-way repeated measures ANOVA was used to analyze changes in behavior among the groups across training. Bonferroni-corrected post hoc comparisons and t tests were made to compare groups on individual sessions. No statistical tests were used to predetermine sample sizes, but our sample sizes were similar to previously published studies. Photometry data were collected with TDT Synapse software and analyzed using MATLAB. All comparisons were two tailed. Data in figures are expressed as the mean ± s.e.m. Statistical significance was set at *p* < 0.05.

### Histology

Rats received i.p. injections of Fatal-Plus (2 ml/kg; Patterson Veterinary) to induce a deep anesthesia, and were transcardially perfused with cold phosphate buffered saline (PBS) followed by 4% paraformaldehyde (PFA). Brains were removed and post-fixed in 4% PFA for ∼24 h, then cryoprotected in a 30% sucrose in PBS for 48 h. Fluorescence from viral injections as well as optic fiber damage location was visualized using standard immunohistochemical approaches. Brain sections were cut at 50 μm on a cryostat (Leica Biosystems). For tyrosine hydroxylase (TH) visualization, we completed immunohistochemistry. Sections were washed in PBS and incubated with bovine serum albumin (BSA) and Triton X-100 (each 0.2%) for 20 min. 10% normal donkey serum (NDS) was added for a 30-min incubation, before primary antibody incubation (rabbit anti-TH, 1:500, Fisher Scientific) overnight at 4 °C in PBS with BSA and Triton X-100 (each 0.2%). Sections were then washed and incubated with 2% NDS in PBS for 10 min and secondary antibody was added (1:200 Alexa Fluor 594 donkey anti-rabbit) for 2 h at room temperature. Sections were washed twice in PBS, mounted on microscope slides, and coverslipped with Vectashield containing DAPI counterstain. Fluorescence from synaptophysin-mRuby and mCherry was not amplified. Slides were imaged using a fluorescent microscope (Keyence BZ-X710) with a 4x and 20x air immersion objective. For photometry and optogenetics experiments, rats were included in VTA or SNc groups only if fiber tips were no more than 500 um dorsal to the target region. Subjects with implants outside of the VTA or SNc were separately analyzed as control/comparison subjects.

